# Functional traits of young seedlings predict trade-offs in seedling performance in three neotropical forests

**DOI:** 10.1101/2023.01.14.523467

**Authors:** Margaret R. Metz, S. Joseph Wright, Jess K. Zimmerman, Andrés Hernandéz, Samuel M. Smith, Nathan G. Swenson, M. Natalia Umaña, L. Renato Valencia, Ina Waring-Enriquez, Mason Wordell, Milton Zambrano R., Nancy C. Garwood

**Affiliations:** Department of Biology, Lewis & Clark College, Portland, OR. USA; Smithsonian Tropical Research Institute, Apartado 0843–03092, Balboa, Republic of Panama; Department of Environmental Sciences, University of Puerto Rico-Río Piedras, San Juan, Puerto Rico. USA; Department of Biology, Colorado State University, Ft. Collins, CO. USA; Department of Biological Sciences, University of Notre Dame, Notre Dame, IN. USA; Ecology and Evolutionary Biology, University of Michigan, Ann Arbor, MI. USA; Facultad de Ciencias Exactas y Naturales Escuela de Ciencias Biológica, Pontificia Universidad Católica del Ecuador, Quito, Ecuador; School of Biological Sciences, Southern Illinois University, Carbondale, IL. USA

**Keywords:** advanced regeneration, Barro Colorado Island, demography, Luquillo, seedling growth, seedling survival, Yasuní

## Abstract

1. Understanding the mechanisms that promote the coexistence of hundreds of species over small areas in tropical forest remains a challenge. Many tropical tree species are presumed to be functionally equivalent shade tolerant species that differ in performance trade-offs between survival in shade and the ability to quickly grow in sunlight.
2. Variation in plant functional traits related to resource acquisition is thought to predict variation in performance among species, perhaps explaining community assembly across habitats with gradients in resource availability. Many studies have found low predictive power, however, when linking trait measurements to species demographic rates.
3. Seedlings face different challenges recruiting on the forest floor and may exhibit different traits and/or performance trade-offs than older individuals face in the eventual adult niche. Seed mass is the typical proxy for seedling success, but species also differ in cotyledon strategy (reserve vs photosynthetic) or other seedling traits. These can cause species with the same average seed mass to have divergent performance in the same habitat.
4. We combined long-term studies of seedling dynamics with functional trait data collected at a standard developmental stage in three diverse neotropical forests to ask whether variation in coordinated suites of traits predicts variation among species in demographic performance.
5. Across hundreds of species in Ecuador, Panama, and Puerto Rico, we found seedlings displayed correlated suites of leaf, stem, and root traits, which strongly correlated with seed mass and cotyledon strategy. Variation among species in seedling functional traits, seed mass, and cotyledon strategy were strong predictors of trade-offs in seedling growth and survival.
6. Our findings highlight the importance of cotyledon strategy in addition to seed mass as a key component of seed and seedling biology. These results also underscore the importance of matching the ontogenetic stage of the trait measurement to the stage of demographic dynamics.
7. *Synthesis:* With strikingly consistent patterns across three tropical forests, we find strong evidence for the promise of functional traits to provide mechanistic links between seedling form and demographic performance.

## Introduction

The coexistence in a single hectare of tropical rainforest of hundreds of tree species has long been a beguiling puzzle for ecologists. Understanding such high levels of local diversity is even more puzzling given that most are functionally equivalent shade-tolerant species (S. J. Wright, 2002). The promise of trait-based ecology is the possibility of predicting species composition through a mechanistic link between the characteristics with which species encounter the environment and the variation in performance among species in that environment (Shipley et al., 2016; Westoby & Wright, 2006; Worthy & Swenson, 2019). Functional traits of plants are morphological and physiological characteristics that reflect constraints in allocation of resources to tissues and structures that affect fitness because they facilitate the acquisition and storage of resources, defense against enemies, and eventually reproduction (Reich et al., 1997, 2003; Westoby, 1998; Westoby et al., 2002). In a spatially and temporally variable environment, trait differences among species should be associated with differences in performance.

Several key axes of trait variation have been reported from a large body of work on tree demography and functional traits. These include the investment in wood per unit volume (i.e. wood density) (Chao et al., 2008; Chave et al., 2009; Enquist et al., 1999; King et al., 2006; Kraft et al., 2010; Lawton, 1984; Putz et al., 1983), investment in leaf biomass or nutrients per unit mass or area (e.g. specific leaf area or leaf nitrogen content) (Kitajima & Poorter, 2010; Poorter et al., 2008; I. J. Wright et al., 2004), and investment in seed provisioning (Baraloto et al., 2005; Muller-Landau, 2010; S. J. Wright et al., 2010). The variation in investment among these strategies results in trade-offs between growth and survival (Adler et al., 2013; Reich et al., 2003). For example, resource investment in higher wood density results in lower average growth rates in girth but reduces damage from physical and biotic factors and increases survival (Brando et al., 2012; King et al., 2006; Russo et al., 2010). Likewise, leaf lifespan and photosynthetic rates are correlated with the rate of leaf production, turnover, and structural quality of leaves (Reich, 2014). Species with high maximum photosynthetic rates, for example, may be able to construct leaves without heavy defense investment because these leaves also may be replaced if lost (Coley et al., 1985; Kitajima, 1994). In this way, short leaf lifespan and high turnover rates are linked to high maximum photosynthetic rates (Reich, 2014; I. J. Wright et al., 2004). The great variation observed among species in seed mass reflects a tradeoff between production of many small or fewer large seeds. Colonization ability is enhanced by seed number while stress tolerance is enhanced by large seed reserves to provision young seedlings (Muller-Landau, 2010; Myers & Kitajima, 2007; Paz & Martinez-Ramos, 2003; Venable, 1996, p. 199; Visser et al., 2016; Westoby, 1998).

Seedlings face different challenges recruiting on the forest floor than older individuals face in the eventual adult niche (Spasojevic et al., 2014). There is increasing empirical evidence that non-random mortality at young seedling stages is an important filter setting the template for forest composition (Green et al., 2014; Zhu et al., 2015). The great majority of species in tropical forests are shade tolerant. Their seedlings must germinate and survive in the shaded understory, where they face asymmetric competition from larger individuals, and must defend against natural enemies, which cause particularly high mortality at these young vulnerable stages (Kitajima & Myers, 2008). The seedling traits that confer success against these challenges are usually represented by seed mass, with the understanding that variation in seed mass in part results from variation in nutrient reserves available to nourish the young plant (Grubb, 1977). Thus, production of fewer large seeds can result in seedlings with more resources to withstand stresses (and increase survival rates), while production of many, smaller seeds can increase rates of dispersal to suitable germination sites at the cost of higher mortality rates in less favorable sites (Muller-Landau, 2010).

An understudied aspect of the demographic consequences of resource provisioning supplied to young seedlings is the type of cotyledon strategy a species possesses (Baraloto & Forget, 2007; Garwood, 1996, 2009; Kitajima, 1996; Kitajima & Myers, 2008; Zanne et al., 2005). Some species have reserve-type cotyledons that store carbohydrates and nutrients. They draw on these reserves to build photosynthetic machinery and other tissues to survive stresses of low light or attack by herbivores and pathogens (Green & Juniper, 2004; Myers & Kitajima, 2007). Other species have leafy, photosynthetic cotyledons that emerge from the seed at germination and must begin carbon acquisition immediately to build plant tissues (Ganade & Westoby, 1999). While species with the reserve cotyledon type tend to have larger seeds than those with photosynthetic cotyledons, seed mass varies by several orders of magnitude within each cotyledon strategy, even among closely related taxa (e.g. Baraloto & Forget, 2007; Kelly, 1995). Though framed here as a cotyledon reserve strategy, we include in this category species that store reserves in the hypocotyl or in the root of the embryo, and those that absorb energy from copious endosperm via specialized haustorial cotyledons (Garwood, 2009).

Here we present longitudinal studies of seedling demography with seedling trait data collected at a standard developmental stage from understory field conditions of three diverse neotropical forests. We compared trait variation among species at each site to the growth and survival of their newly recruited seedlings. Specifically, we asked:

i. How do species vary in leaf, stem, or root traits at the young seedling stages?
ii. Is variation in leaf, stem, or root traits related to variation in seed mass and/or cotyledon strategy among species?
iii. Is there a demographic trade-off among species between growth rates and survival rates of newly germinated seedlings?
iv. Do differences in leaf, stem, and root traits and/or cotyledon strategy or seed mass predict demographic trade-offs?

We hypothesize that a species’ cotyledon type is an important indicator of a species’ investment strategy, and thus should predict differences in both functional traits and the fitness consequences of those traits. The species with seed storage reserves can afford greater investment in tissue density or chemical composition as a strategy for withstanding the stresses of the understory, but this investment would come at a cost of reduced growth rates. The species with leafy photosynthetic cotyledons and no storage reserves must instead invest in resource acquisition, resulting in higher growth rates when light is available, but also higher mortality when light is low due to a lack of stored reserves.

## Materials and Methods

### Research sites

Since 2002 in Yasuní National Park, Ecuador, since 1994 at Barro Colorado Island (BCI), Panama, and since 2007 in Luquillo, Puerto Rico, we have been monitoring seedling dynamics within large (16-50 ha) forest dynamics plots (FDPs). The Pontificia Universidad Católica del Ecuador (PUCE) manages the Yasuní Scientific Station (ECY) in Yasuní National Park, Ecuador (0° 41’ S, 76° 24’ W). This site has relatively constant temperatures and an everwet climate, with no month receiving less than 100 mm of rainfall on average and annual rainfall typically exceeding 3000 mm (Valencia et al., 2004). The park, and the adjacent Huaorani Ethnic Reserve, comprise the largest protected area in Ecuador. The 50-ha FDP contains more than 1100 tree species, making it one of the most diverse forests ever studied (Losos & Leigh, 2004).

Robin Foster and Stephen Hubbell established the 50-ha FDP on BCI (9° 9’ N, 79° 51’ W) in 1982. Monthly temperatures average 26° C for 11 months and 27° C in April. Annual rainfall averages 2,600 mm, with 10% of the annual total falling during a four-month dry season (usually mid-December into April). The 50-ha FDP contains 300 species of trees in old-growth forest, which escaped human disturbance for the past 500 years and never supported agriculture (Piperno, 1990).

The Luquillo 16-ha FDP (18° 9’ N, 65° 49’ W) was established in 1990 near the El Verde Field Station of the Luquillo Experimental Forest in Puerto Rico. The everwet site receives an average of 3500 mm of rainfall per year, with no month averaging <200 mm. Luquillo has experienced recent anthropogenic and natural disturbances (Thompson et al., 2002). Before 1934, the northern part of the plot was used for agriculture while the southern third of the plot was selectively logged (Zimmerman et al., 1994). Additionally, the forest is periodically damaged by hurricanes, with two major hurricanes in the period that is relevant for this study (hurricane Hugo in 1989, hurricane Georges in 1998) (Zimmerman et al., 2018).

### Seedling demography

Within each FDP, we established a network of seedling plots where we conduct annual surveys of seedling dynamics in association with long-term monitoring of flowering and fruiting phenology (Metz, 2012; S. J. Wright et al., 2016), using standardized methodology that facilitates comparisons among sites (Anderson-Teixeira et al., 2015; Metz, 2007). One raised mesh seed trap surrounded by three 1 m^2^ seedling plots comprises a census station, for a total of 600 plots (in 200 stations) at Yasuní, 800 plots in 250 stations at BCI including 50 plots originally located in natural treefall gaps, and 360 plots in 120 stations at Luquillo. Most plots are in the shaded understory, although treefall gaps create temporarily open canopy over some plots.

Each year in each plot, we tag and measure all seedlings of woody species (including trees, shrubs, palms, and lianas), identify all newly recruited seedlings, and determine the fate of previously tagged seedlings. This allows us to generate species- and age-specific rates of recruitment, growth, and survival. Here the term seedling indicates young plants that have germinated from seeds that have fallen into our study plots. Newly germinated seedlings range in height from a few millimeters to tens of centimeters. There is no lower size or age limit on when a seedling enters our study.

Absolute seedling age for these new recruits depends on when peak germination occurred relative to the timing of the annual census. At Yasuní, seed germination occurs throughout the year (Persson, 2005) but the annual seedling census occurs in May-July. Newly recruited seedlings thus range in age from 1 to 364 days (e.g., the potential range of days between the unknown date of germination and the annual census). At Luquillo, seedling ages are similar because germination also occurs throughout the year, and the annual seedling census occurs March-May. At BCI, seed germination is more synchronized:, most species germinate in May and June but some as late as November (Garwood, 1983). The annual seedling census occurs in January–March, thus newly recruited seedlings are at least two months post-germination when first censused, and most are 7-10 months old.

We analyzed growth and survival between the first and second censuses for each seedling for 15 cohorts at Yasuní, 21 cohorts at BCI, and eight cohorts at Luquillo. These seedlings recruited between 2003-2018 at Yasuní, 1995-2015 at BCI, and 2008-2015 at Luquillo (after baseline censuses in 2002, 1994, and 2007, respectively). We limited our analyses to species that had seedling recruits in ≥ 5 years and at least 5% of census stations (i.e., ≥ 10 census stations for Yasuní and BCI and ≥ 6 stations for Luquillo) across the study period and at least 8 surviving individuals to calculate growth rates. This included 27,747 individuals from 216 species at Yasuní, 67,205 individuals from 188 species at BCI, and 45,117 individuals from 43 species at Luquillo. The sample size for each species ranged from 10-1446 individuals at Yasuní (median=52.5; mean=128.5), from 13-9052 individuals at BCI (median=119; mean=357.5), and from 14-11802 individuals at Luquillo (median=133, mean=1049.2).

We scored whether each individual seedling survived the first year after entering our censuses and pooled individuals across census cohorts to quantify the study-wide first-year survival proportion for each species. We calculated a relative growth rate over the first year for surviving individuals (ln*(ht_1_)-*ln*(ht_0_)*)/((*date_1_*-*date_0_*)/365.25) using the difference between initial recruitment height (*ht_0_*) in the first census in which a seedling appeared and its height the following year (*ht_1_*) and standardizing to one year. We then calculated species-specific average growth rates using the mean growth rate of individuals in each census plot, to account for their non-independence, before taking the mean across plots for the species rate.

### Seedling functional traits

We assessed a suite of functional traits on young, field-collected seedlings. At Yasuní, these were individuals at a standardized developmental stage, when their first photosynthetic organ (whether cotyledon or leaf) was fully expanded and matured but before further shoot growth (Alvarez-Clare & Kitajima, 2007; Kitajima, 2011; Myers & Kitajima, 2007). We measured traits on 747 field-harvested individuals from 138 species collected in and around the Yasuní FDP. Replicate sample sizes ranged from 1-13 (median=6), and only 22 species had three or fewer replicate individuals. Because species germinate throughout the year, have different lag periods between seed fall and seed germination, differ in recruitment densities among years, and grow at different rates, the seedlings used in this study were collected across several years, each having reached this standard developmental stage at different points in the calendar year.

At BCI, we harvested 484 newly recruited seedlings from the forest between 29 January and 10 June 2015. This included 1-6 replicates (median=4) for each of 145 species. These seedlings were likely six to 10 months old when harvested because the majority of seedlings germinate in May and June (Garwood, 1983). Previous studies at BCI have shown traits at this stage (several months after leaf expansion) are well correlated with the traits measured one month after expansion of the first photosynthetic organ (Alvarez-Clare & Kitajima, 2007).

Seedlings (<50 cm in height) at Luquillo were harvested from a location near the forest dynamics plot in 2014 between June and July. We collected seedlings of 50 species; sample size varied 1-357 with seven species represented by a single individual.

After harvesting, we scored the cotyledon type of every species using the categories of Garwood (2009). We divided cotyledon types into two broad classes – those with photosynthetic cotyledons versus non-photosynthetic storage cotyledons.

At Yasuní, we scanned freshly harvested seedlings at high resolution and used ImageJ64 (Rasband, 2018) to assess the area of photosynthetic leaves and/or cotyledons, stem length from the apical meristem to root collar, and length of the longest root from root collar to tip. Seedlings were then dried at 65°C before we weighed the mass of all parts separately (roots, stem, leaves and cotyledons with and without petioles, and any other seedling tissues).

At BCI, we recorded the number of cotyledons and leaves, measured plant height (from root collar to tallest meristem) and length of the longest root (from root collar to root tip) with a ruler, and scanned each freshly harvested seedling. For each seedling, we determined fresh mass of stem, roots, and pooled petioles, and fresh mass and area (Licor 3100, Lincoln, Nebraska) of cotyledons, pooled leaf laminas and two individual leaf laminas. For stems, we recorded length, diameter at base and tip, and volume by water displacement. Finally, we recorded dry mass after drying at 60°C of the stem, roots, cotyledons, pooled petioles, pooled laminas and two individual laminas for each seedling.

Seedlings at Luquillo were dried and weighed to determine stem, leaf, and root dry mass. We then selected 1-3 healthy and fully developed leaves per seedling to scan and measure SLA. The scanned leaves were then dried and weighed separately. Further details on the trait data collection at Luquillo can be found in Umaña et al. (2015).

We used these masses, areas, and lengths to calculate seedling leaf, stem, and root traits at each site (Cornelissen et al., 2003; Pérez-Harguindeguy et al., 2016). We calculated specific leaf area (SLA) by dividing the total area of photosynthetic material (leaves and/or cotyledons, without petioles) by the dry mass of the same material (Cornelissen et al., 2003). We estimated fresh stem volume using the length of the stem, two diameter measurements and the equation for volume of an elliptical cylinder. In rare cases, stems were rectangular, and we adjusted the calculation of volume accordingly. We calculated specific stem density (SSD) by dividing stem dry mass by fresh stem volume (Cornelissen et al., 2003). We calculated a modified version of specific root length (SRL) appropriate to seedling structure by dividing the length of the longest root by the total dried root mass (Cornelissen et al., 2003; Kramer-Walter et al., 2016). Finally, we calculated the root to shoot biomass ratio (RS) by dividing root dry mass by the dry mass of all aboveground parts (stem, leaf, and cotyledons including petioles). Trait values were averaged over replicate individuals to obtain species-level trait estimates (Suppl. Table 1).

We measured mean dry seed mass from a pooled sample of seeds (1 to >200, depending on seed size) from one or several individuals where seeds were collected from fruiting individuals or fell into seed traps from our long-term studies of fruiting phenology at Yasuní. We measured dry seed mass for up to five seeds from up to five fruits from up to five individuals of each species at BCI (S. J. Wright et al., 2010). Seed masses at Luquillo were measured on at least three individual dried seeds collected from the seed traps arrayed in the Luquillo forest dynamics plot (Swenson et al., 2012). Here, we use diaspore dry mass, as a proxy for true seed mass, where the diaspore includes the endocarp and testa and, for wind-dispersed species, any appendages that facilitate dispersal.

### Statistical analyses

We asked whether species varied in suites of seedling functional traits and whether seed mass and/or cotyledon strategy were useful indicators of the trait associations. We examined variation in seed mass and cotyledon strategy using t-tests comparing species with reserve- and photosynthetic-type cotyledons. We explored coordinated variation in leaf, stem, and root functional traits (i.e., to identify suites of traits), using nonmetric multidimensional scaling (NMDS) analyses across 138 species at Yasuní and 143 species at BCI with measures of specific leaf area (SLA), specific stem density (SSD), specific root length (SRL), and root-to-shoot biomass ratio (RS). We next asked whether seed mass, cotyledon type (photosynthetic vs. reserve), and/or their interaction predicted significant differences in the variation of SLA, SSD, SRL and RS using a permutational multivariate analysis of variance (ADONIS) for the subset of 113 species at Yasuní and 128 species at BCI with trait and seed mass data. We conducted both analyses in the R package *vegan* (Oksanen et al., 2018).

To determine whether first year seedling dynamics reflect a trade-off between growth and survival, we conducted a standard major axis regression (SMA) using the *smatr* R package (Warton et al., 2012) after first logit-transforming survival proportions (Warton & Hui, 2011) and log-transforming relative growth rates. We conducted regressions individually for each site using species with reserve and photosynthetic cotyledons and additional species with cotyledon type not yet classified. We also pooled all species with reserve or photosynthetic cotyledons from all three sites into one SMA analysis and tested for a shift along the axis between cotyledon types or whether the slopes differed among sites. Finally, after understanding demographic trade-offs and links between seed mass and the other traits, we created two composite variables using the R package *vegan* (Oksanen et al., 2018) and asked whether variation in seed mass and seedling traits predicted interspecific variation in seedling demography. We standardized the species survival and relative growth rate variables described above (mean=0, standard deviation=1) and reduced the growth-survival trade-off continuum to a single axis using a principal components analysis (PCA), which we refer to here as our *demographic PCA*. The first axis of this demographic PCA accounted for 70.5% of variation among species at Yasuní, 75.6% at BCI, and 66.1% at Luquillo. We used first axis demographic PCA scores to represent the location of each species on the continuum between high survival and slow growth versus lower survival and faster growth. Next, we reduced SLA, SSD, SRL, RS, and seed mass to orthogonal composite variables using PCA, with data first log-transformed and standardized. The first axis of this *trait PCA* explained 48.2% of the variation in traits at Yasuní and 45.8% at BCI, and for each site, the first PCA axis was most strongly positively associated with seed mass and negatively associated with specific leaf area and specific root length (Suppl. Table 4). Though fewer traits were available for seedlings at Luquillo, we similarly used PCA to reduce SLA, RS, and seed mass to one axis explaining 49.9% of variation in these traits. We used the first trait PCA axis to represent a species’ position in a continuum of variation in coordinated suites of functional traits.

We then conducted an analysis of covariance of species’ position on the first demographic PCA axis (a measure of the growth-survival trade-off continuum) as the dependent variable against the independent variables of cotyledon type crossed with species’ scores on the first trait PCA axes. This analysis included the 63 species at Yasuní, 92 species at BCI, and 28 species at Luquillo for which we had all traits, seed mass, and seedling performance data with the sample size restrictions described above. All analyses were conducted in the R statistical programming language version 3.5.0 (R Core Team, 2018).

## Results

Species with reserve-type cotyledons at all sites had significantly higher seed mass than those with photosynthetic cotyledons (Yasuní: t=5.10, p<0.0000021; BCI: t=7.32, p<5.4×10^-11^; and Luquillo: t=4.433, p<7.66×10^-5^), but there was much overlap in seed mass between cotyledon types at all sites (Fig. 1).

**Figure 1.**
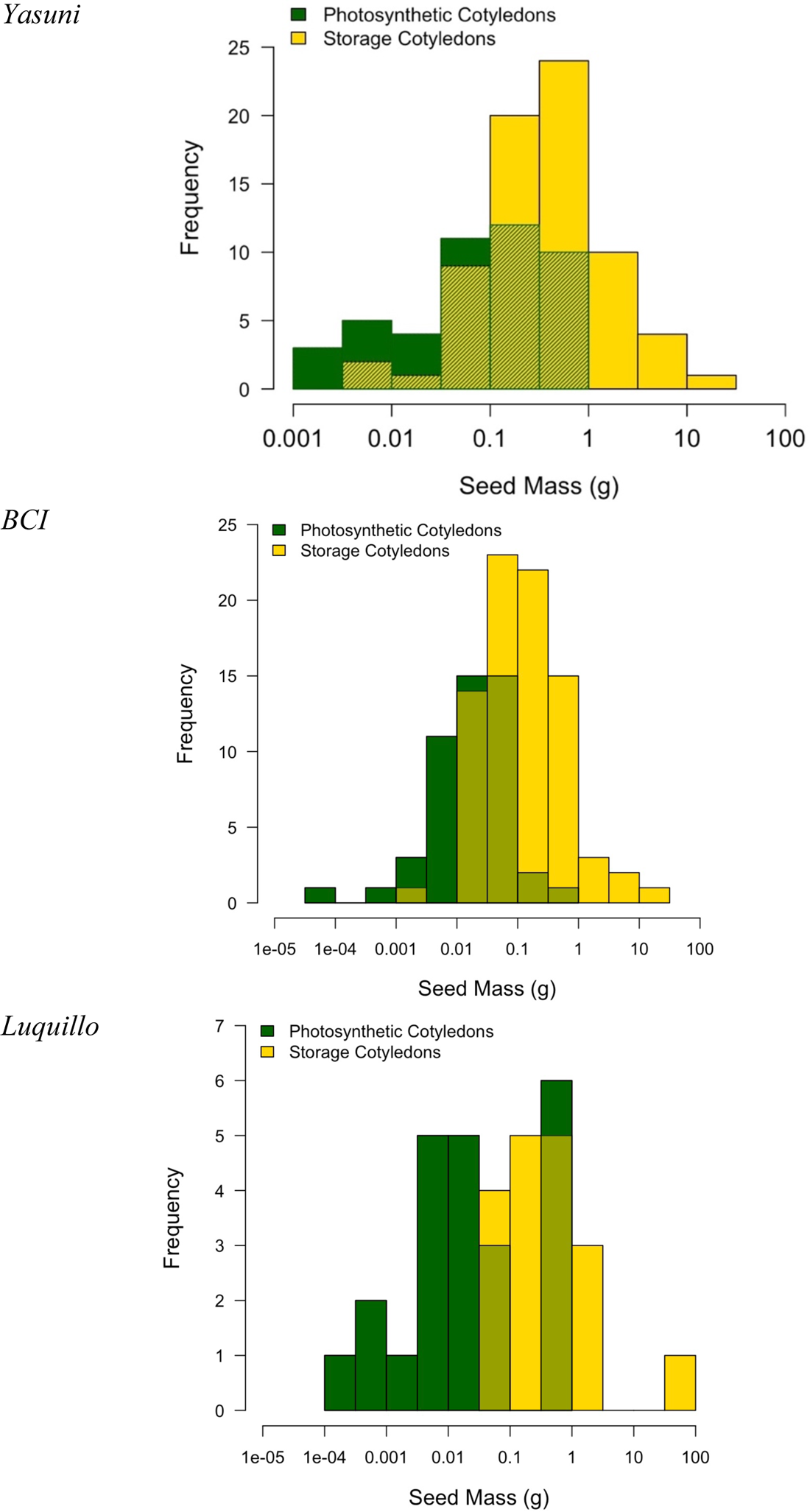
Seed mass variation by cotyledon type for 116 species at Yasuní, 130 species at BCI, and 41 species at Luquillo. Green bars are for species with photosynthetic cotyledons and yellow bars indicate species with storage cotyledons. The light green shading shows the overlap in the histograms of the two groups.

There were similarities across sites in the relationships among individual traits, and much overlap in trait range values between cotyledon strategies (Suppl. Tables 1-2). Seed mass was strongly negatively correlated to specific leaf area at all three sites and to specific root length at BCI and Yasuní (Suppl. Table 2). In contrast, seed mass was poorly correlated with stem density or root:shoot ratio, except at Luquillo. The PCA loadings for each of these traits were also remarkably similar between Yasuní and BCI (Suppl. Table 4).

**Table 1.**
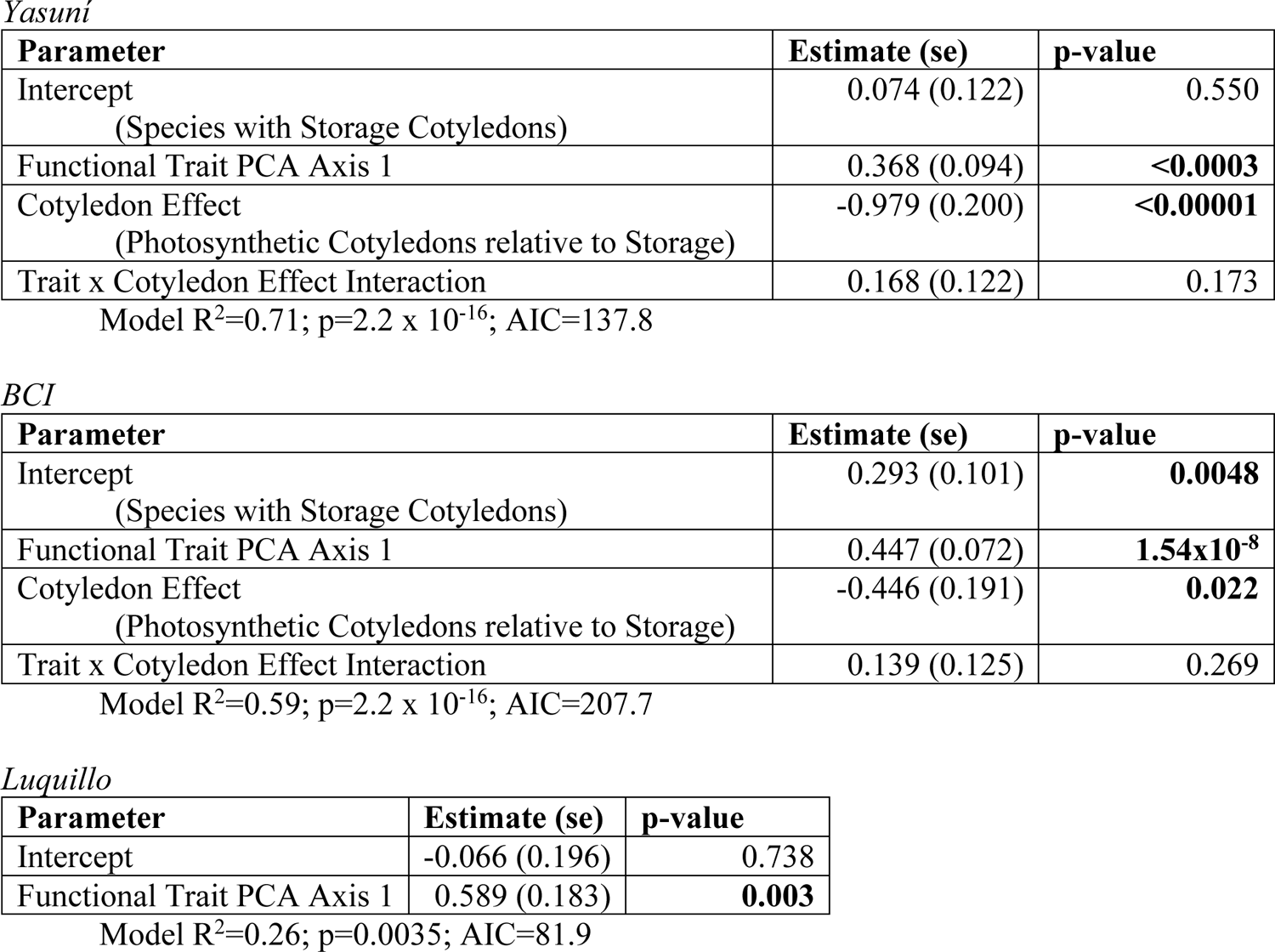
Regression coefficients for model of position along the growth-survival trade-off axis (dependent variable) against the composite functional trait axis crossed with cotyledon strategies (independent variables). Here the trait PCA axis includes seed mass in the composite variable. Positive slope estimates indicate a positive relationship with seedling survival and a negative relationship with seedling growth in the first year after recruitment. The Luquillo analysis contains only the trait axis as a predictor because adding the additional parameters of cotyledon strategy and its interaction with seed mass increased the model AIC.

*Yasuni*
| Parameter | Estimate (se) | p-value |
| --- | --- | --- |
| Intercept<br>(Species with Storage Cotyledons) | 0.074 (0.122) | 0.550 |
| Functional Trait PCA Axis 1 | 0.368 (0.094) | **<0.0003** |
| Cotyledon Effect<br>(Photosynthetic Cotyledons relative to Storage) | -0.979 (0.200) | **<0.00001** |
| Trait x Cotyledon Effect Interaction | 0.168 (0.122) | 0.173 |

*BCI*
| Parameter | Estimate (se) | p-value |
| --- | --- | --- |
| Intercept<br>(Species with Storage Cotyledons) | 0.293 (0.101) | **0.0048** |
| Functional Trait PCA Axis 1 | 0.447 (0.072) | **$1.54 \times 10^{-8}$** |
| Cotyledon Effect<br>(Photosynthetic Cotyledons relative to Storage) | -0.446 (0.191) | **0.022** |
| Trait x Cotyledon Effect Interaction | 0.139 (0.125) | 0.269 |

*Luquillo*
| Parameter | Estimate (se) | p-value |
| --- | --- | --- |
| Intercept | -0.066 (0.196) | 0.738 |
| Functional Trait PCA Axis 1 | 0.589 (0.183) | **0.003** |

Model $R^2=0.26$ ; $p=0.0035$ ; AIC=81.9

Together, seed mass, cotyledon type, and their interaction explained 52.2% and 39.5% of the variation in multivariate trait space for SLA, SRL, SSD and R:S at Yasuní and BCI, respectively (Suppl. Table 3). The significant interaction between seed mass and cotyledon type indicates an important role for early resource acquisition strategies (reserve vs. photosynthetic cotyledons), no matter the seed mass. Species with storage reserve cotyledons tended to invest more mass per unit volume of stem or root, and generally grew from seeds with larger mass (Fig. 2). There was a wide range of variation in traits and seed mass among species with either cotyledon strategy (Fig. 2).

**Figure 2.**
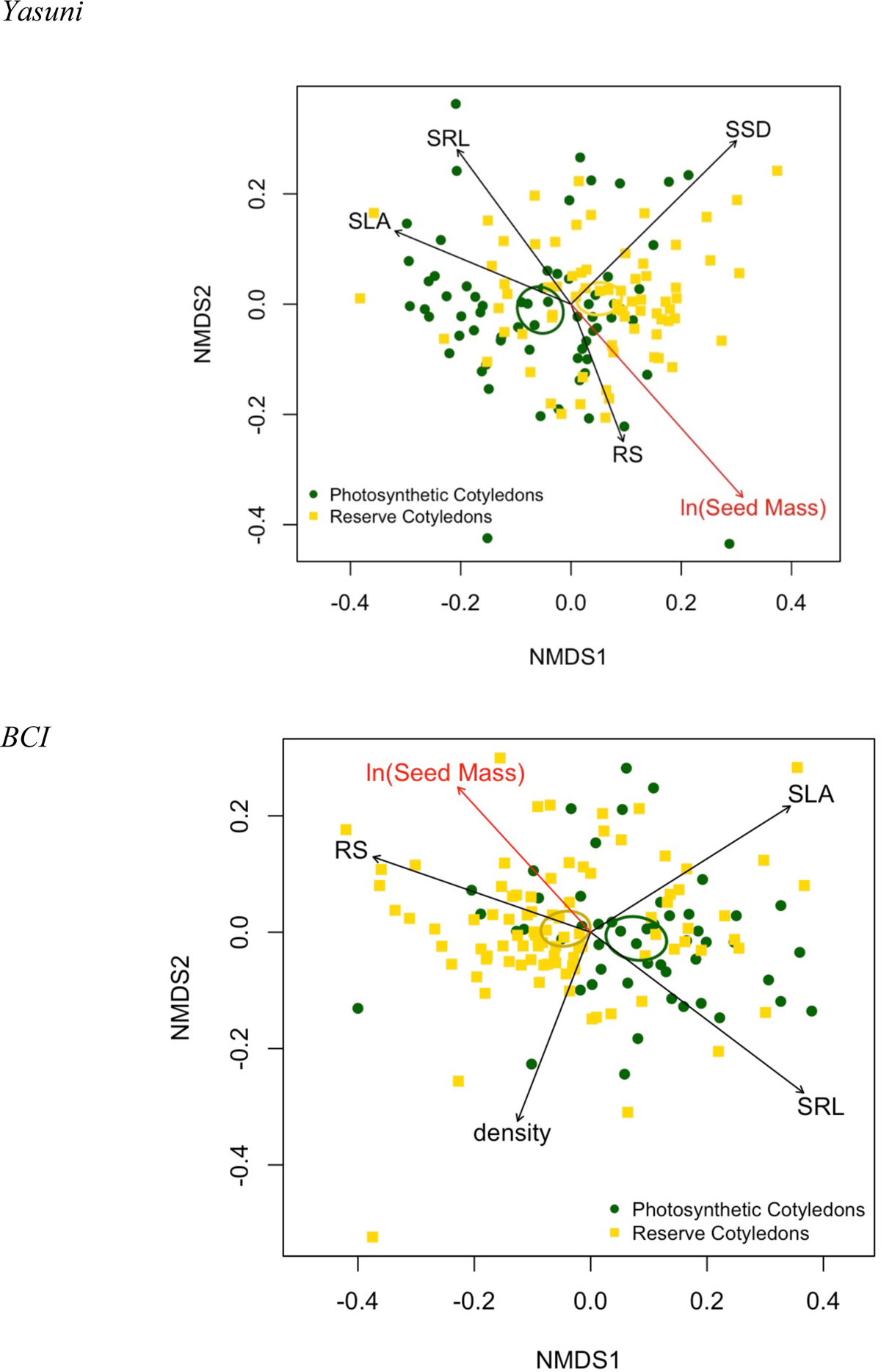
NMDS of four functional trait values (specific leaf area, stem density, specific root length. root:shoot biomass ratio). Black arrows indicate strength and direction of maximum correlation of the trait with the ordination space. Seed mass, which was not used to create the ordination, was significantly correlated (p<0.001) with the ordination space and is overlaid with a red arrow. Green and yellow symbols represent species with photosynthetic and reserve cotyledons, respectively.

There was a strong tradeoff in the survival and growth of newly recruited seedlings at all three sites (Fig. 3) and separately at Yasuní and BCI (Yasuní: R^2^=0.169, p < 3.4×10^-10^; BCI: R^2^=0.26, p < 5.5×10^-14^) with a weaker but similar trend at Luquillo (R^2^=0.10, p<0.035) (Suppl. Fig. 1). High rates of survival in the first year occurred in species with slower relative growth rates (Fig. 3). Cotyledon type, seed mass and seedling traits (SLA, SRL, SSD and RS) combined to explain 69% at Yasuní and 59% at BCI of the interspecific variation in species’ positions on the growth-survival tradeoff continuum (Table 1). At Yasuní, including cotyledon strategy as a predictor explained significantly more of the variance in the demographic PCA (the R^2^ increased from 0.59 in the model without cotyledon strategy to 0.71 in the full model) (Suppl. Table 5). At Luquillo the lowest AIC model contained only the combined seed mass and seedling trait PCA axis (across fewer traits than the other two sites) and explained 29% of variation in the growth-survival tradeoff. Across all species at both sites, those with lower SLA, lower SRL, and greater seed masses experienced greater survival at a cost of slower relative growth rates (Fig. 4). Even with similar suites of functional trait and seed mass values, seedlings with reserve cotyledons tended towards higher survival and lower relative growth rates relative to species with photosynthetic cotyledons (Fig. 4; Table 1).

**Figure 3.**
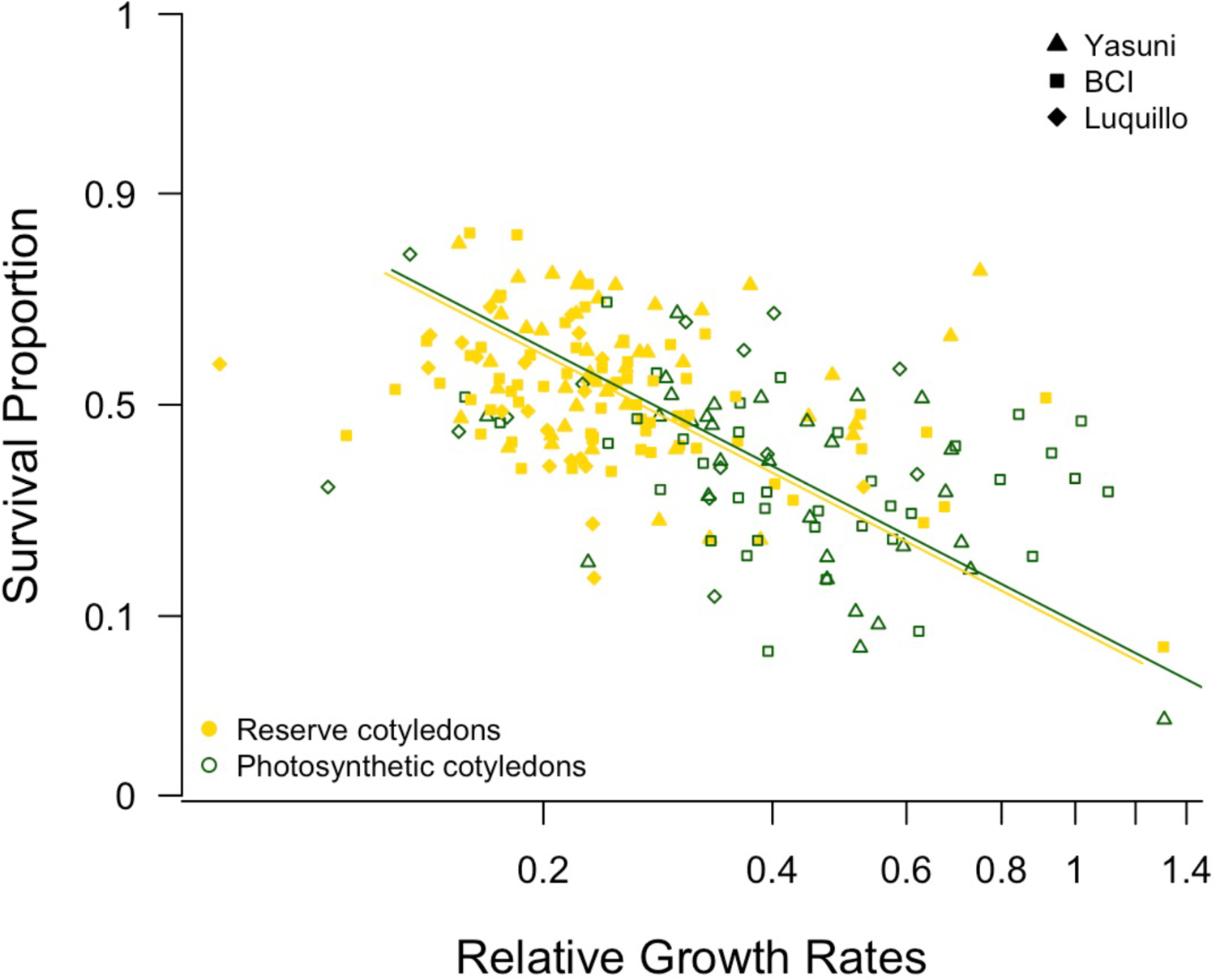
Single-major axis regression of logit-transformed proportion of first-year seedling survival and log-transformed average relative growth rate of surviving seedlings. Green and yellow symbols represent species with photosynthetic versus reserve cotyledons, respectively, with different shapes indicating the study site.

**Figure 4.**
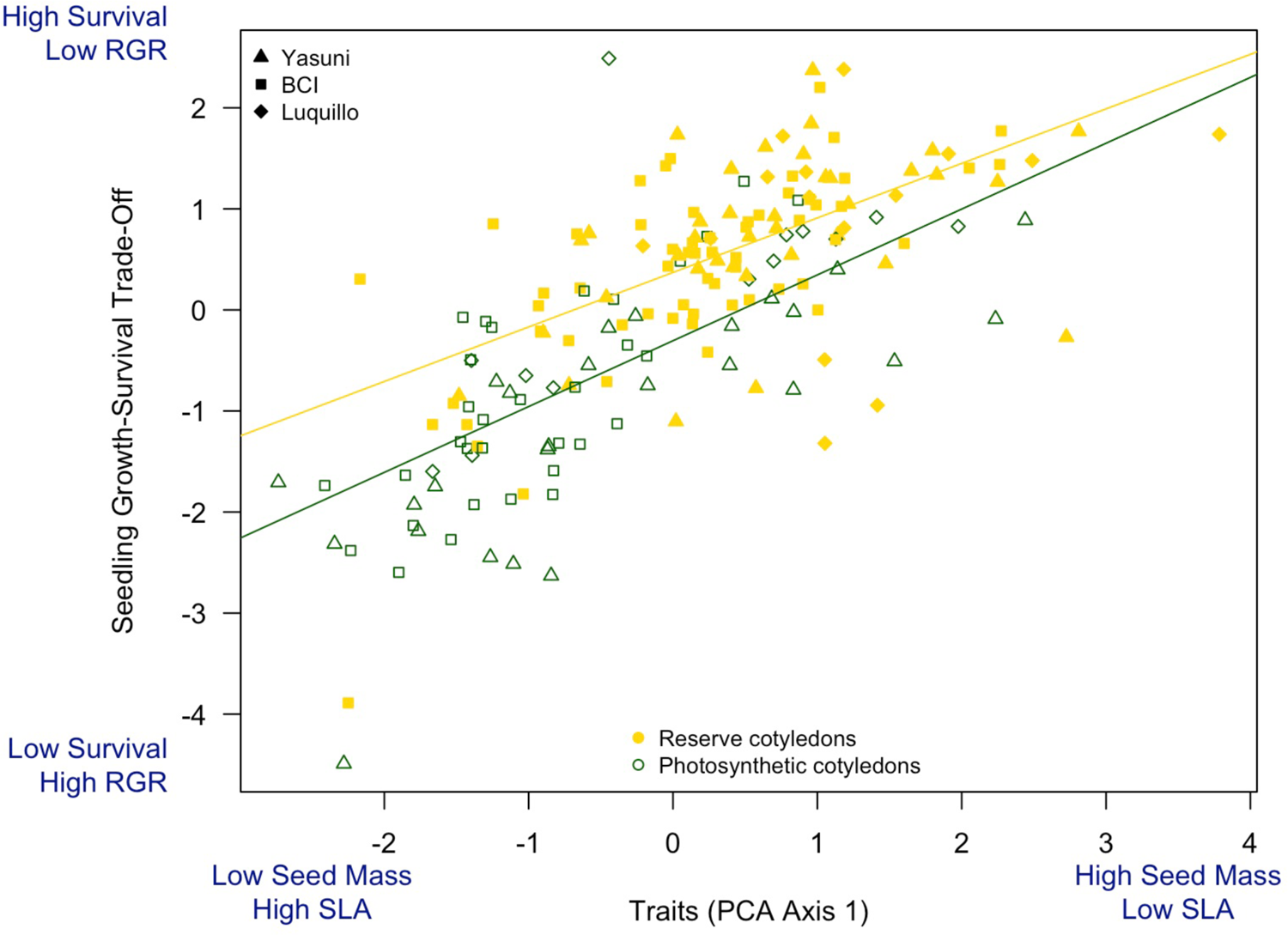
Variation in seedling functional traits (of leaves, stems, roots and seeds) in combination with cotyledon type explains much of the variation in species’ position along a continuum of high survival and high relative growth.

## Discussion

We have demonstrated that interspecific variation in seedling functional traits, seed mass, and cotyledon strategy are strong predictors of trade-offs in seedling growth and survival for hundreds of woody species from three neotropical forests. First-year seedlings displayed correlated leaf, stem, and root traits (Table 1), and these trait suites were strongly associated with interspecific variation in seed mass and cotyledon strategy (Table 2). These results are intriguing because of the striking similarities in the role of traits in predicting demographic tradeoffs across several hundred species from the Western Amazon, southern Mesoamerica and the Greater Antilles (Table 3).

The similarities across sites in this study contrast with other multi-site studies suggesting little to no demographic predictive power of functional traits (i.e. low R^2^ values such as, for example, 0.03 (Paine et al., 2015), 0.39-0.43 (S. J. Wright et al., 2010), and others reviewed by Worthy and Swenson (2019). We found much greater explanatory power (i.e. R^2^ of 0.71 at Yasuní, 0.59 at BCI, and 0.26 at Luquillo, which had fewer functional traits available for the analysis, Table 3). Still other studies have urged caution about broad conclusions regarding traits and demography because the importance of a particular trait can change dramatically with the ontogenetic stage examined (Gibert et al., 2016; Visser et al., 2016) or because relationships among traits are weak when measured under field conditions (Laughlin et al., 2017). In some cases where little predictive power of traits on demography has been found, traits were gathered from databases or approximated from genus-level averages (Paine et al., 2015), which may have been measured in a very different environmental context. Our strongly predictive findings argue for the importance of measuring traits at the relevant ontogenetic stage and the relevant environmental context to predict species differences in performance.

Our results corroborated the use of seed mass as a proxy for seedling strategies. Seed mass was highly negatively correlated with two seedling traits (SLA and SRL) that accounted for much of the trait variation we measured, as has been shown elsewhere (Bergmann et al., 2017; I. J. Wright & Westoby, 1999). SLA and SRL quantify investment per unit area or unit length for leaves and roots, respectively. Here, species with larger seeds had thicker or denser leaves and roots (or lower SLA or SRL, respectively). Species with thicker tissues had greater survival and slower growth. This is consistent with findings elsewhere that link resource allocation to mechanisms of survival, for example that finer roots have been shown to be more susceptible to pathogens (Bergmann et al., 2017). These findings also align with evidence that larger seeds contain more energy and nutrient reserves to nourish the developing seedling (Kitajima, 2011), and with theoretical expectations that seed size indicates a trade-off of fecundity vs. stress tolerance for greater survival in resource-poor environments (Muller-Landau, 2010).

Our findings further highlight the importance of cotyledon strategy as a key component of seed and seedling biology. Both large seed size and storage cotyledon strategies have been implicated in increased survival and resprouting following seedling herbivory (Armstrong & Westoby, 1993; Harms & Dalling, 1997). At our sites, species with photosynthetic cotyledons tended to have lower survival and higher relative growth than species with similar functional traits and seed mass but storage-type cotyledons (Fig. 4). Reserve cotyledons confer a survival advantage, no matter the seed mass or values of other leaf and stem traits, because the young plant can draw on the reserves in periods of low light availability (Boot, 1996) or after tissue loss caused by herbivores or other agents (Baraloto & Forget, 2007; Harms & Dalling, 1997). A disadvantage of reserve cotyledons is that the first photosynthetic organs must be built from these reserves following germination (Kitajima & Fenner, 2000), so these species will be slow to capture ephemeral pulses of high light availability in the otherwise relatively dark understory. By contrast, the leafy cotyledons that emerge from a germinating seed of a species with photosynthetic cotyledons can photosynthesize immediately (Kitajima & Fenner, 2000). However, these species have fewer reserves to sustain survival through stressful periods. Both cotyledon strategies occur across orders of magnitude of seed mass variation (Suppl. Fig. 1) and represent a key strategy difference in initial abilities to acquire carbon no matter the nutrient provision indicated by seed size.

In adult trees, differences in SLA are linked to variation in photosynthetic rates and the resource costs of replacing lost or damaged leaves (Reich et al., 1991). High SLA leaves are thought to be more “cheaply” constructed and easy to replace given adequate resources (Worthy & Swenson, 2019; I. J. Wright et al., 2004). Kitajima et al. (2013) measured leaf lifespan for seedlings of many of the BCI species reported here and found that lifespan correlated positively with tissue density. All our traits were measured on seedlings at developmental stages where the first photosynthetic organ (whether leaves or cotyledons) had fully matured (at Yasuní), or when 1-2 sets of true leaves had developed (at BCI or Luquillo). Higher SLA may indicate a shorter leaf lifespan if the relative thinness of leaf tissues makes these leaves more susceptible to insect attack or physical damage (Alvarez-Clare & Kitajima, 2007; Kitajima et al., 2013), however other traits such as the cellulose content of leaves contribute to leaf toughness and are independent of leaf thickness (Westbrook et al., 2011). At these early life stages, loss of the only photosynthetic organs would be a serious threat to survival, which may explain the strong link between a species’ investment in thick tissues and its position along a continuum of high survival versus high relative growth.

Our study design has several strengths that likely contribute to the strong predictive power we found between functional traits and seedling demography in these forests. First, as individuals age and grow, their individual histories (e.g. recent tree gap opening nearby or long-term attack by pathogens) cause much variation in survival and growth that can compound with time. By examining young seedlings, we have eliminated variation in performance that would result from these unknown histories, allowing the signal of functional trait strategies to be more apparent.

Second, we used long-term studies at each site to document growth and survival across forest environments that vary spatially and temporally (due to topography, canopy disturbance, interannual variation in climate, etc.). Many trait studies examine only static snapshots of plant composition or demographic changes over short time intervals. We examined first-year seedling survivorship and growth averaged over 8-21 years, so our data include individuals of a given species responding to a wide range of environmental conditions. Functional trait strategies that have evolved over much longer time scales may not be optimized to the conditions present in a short time interval, which could contribute to year-to-year variability in performance, despite being successful or advantageous in the long-term.

Investigations of these key fitness traits and resulting performance differences among species have often focused on the life histories of the adult plant, such as wood anatomy and growth rate differences between lianas and trees, growth and survival trade-offs in pioneer species vs. later successional trees, or the different environments faced by understory trees vs. canopy emergent (S. J. Wright et al., 2010). Many traits change with position in the tree (e.g. sun vs. shade leaves), but there are few studies on how traits themselves change through ontogeny and whether those differences affect the predictive ability of those traits for other stages. Some recent studies have examined how the relative importance of different traits changes through ontogeny (Gilbert et al., 2006; Hérault et al., 2011; Lasky et al., 2015; Visser et al., 2016), but the plant traits used in these studies have typically been measured on older life stages. For example, Visser et al. (2016) used seed mass and adult traits as predictors of growth and survival across multiple life stages at BCI and found much less explanatory power (full model R^2^=0.10 for seedling growth and 0.25 for seedling survival) than our BCI model using traits collected on seedlings, where traits explained >59% of the variation in the species growth and survival. Although these models were constructed differently and may not be directly comparable, the difference demonstrate the important fact that young seedling traits may not correlate with those that predict success in the adult niche, even though these adult traits, e.g. wood density and specific leaf area, are often used to predict seedling demography.

The strikingly consistent findings across sites in the Western Amazon, southern Mesoamerica and the Greater Antilles in which functional traits account for substantial variation in demographic performance is strong evidence for the promise of traits to provide a mechanistic link between form and function. The abundance of studies that demonstrate low explanatory power of traits in sapling or adult demography must be considered in light of the large variation introduced from unknown individual histories over the many years needed to reach a particular size class. Further we urge caution regarding the frequency with which trait values are obtained from databases, species or individuals at different sites or in different environmental conditions, and, importantly, at ontogenetic stages that do not match the demographic dynamics being examined. Our findings contribute to a growing understanding that the environmental context of the trait is crucial to understanding its role in demography (Swenson et al., 2020; Yang et al., 2018).

## Acknowledgements

We thank the people and government of Ecuador, Panama, and Puerto Rico for protecting their forests and making them available for study. We thank the Ecuadorian Ministerio del Ambiente for permission to work in Yasuní National Park (under N° 014-2019-IC-PNY-DPAO/AVS, N° 012-2018-IC-PNY-DPAO/AVS, N° 008-2017-IC-PNY-DPAO/AVS, N°. 012–2016-IC-FAU-FLO-DPAO-PNY, N°. 014-2015-FLO-MAE-DPAO-PNY, and earlier permits). Completion of the seedling dynamics studies at Yasuní would not have been possible without funding from many generous sources, including the U.S. National Science Foundation (DDIG DEB-0407956; LTREB DBI-0614525, DBI-1122634, and DBI-1754632, DEB-1754668, and REU supplements) and the Center for Tropical Forest Science of the Smithsonian Tropical Research Institute (STRI). For assistance with logistics and data collection on the Yasuní seedling project, we thank A. Loor, E. Zambrano, J. Suarez, P. Alvia, J. Guerrero, L. Shupert, and G. Stonehouse. We thank K. Kitajima for sharing then unpublished seedling trait protocols she developed at BCI. Additional support provided at Luquillo from NSF grants DEB-0620910, 0963447, 129764, and 1546686 to the University of Puerto Rico as part of the Luquillo Long-Term Ecological Research Program. The U.S. Forest Service (Dept. of Agriculture) and the University of Puerto Rico gave additional support. We thank Christopher Nytch, Jimena Forero, and Seth Rifkin for assistance with logistics and data collection, as well as many volunteer field assistants. The manuscript benefited greatly from conversations with P. V. A. Fine and S. A. Queenborough. The authors have no conflicts of interest to declare.

## Author Contributions

Margaret Metz conceived the idea of analyzing relationships between seedling demography and seedling traits data at Yasuní. S. Joseph Wright encouraged the expansion of the study to include Luquillo and BCI. At Yasuní, Metz designed the seedling demography study, Metz and Nancy Garwood designed the seedling trait study, and Metz, Garwood, Renato Valencia, and Milton Zambrano collected data and specimens for seedling traits. Ina Waring-Enriquez, Samuel Smith, and Mason Wordell collected seedling functional trait data. At Luquillo, Jess Zimmerman, Nathan Swenson, and M. Natalia Umaña designed and/or collected data for the seedling demography and traits study. At BCI, Wright and Andrés Hernandéz designed and/or collected data for the seedling demography and traits study. Margaret Metz analyzed the data from the three sites and drafted the manuscript. All authors contributed critically to the drafts and gave final approval for publication.

## Data Availability Statement

Yasuní seedling dynamics data are in preparation and review for storage at the Environmental Data Initiative (EDI) repository, where we will also place the functional trait data. Luquillo seedling dynamics data are already available at EDI through the LTER program at that site. Luquillo seed mass and seedling trait data are already deposited in Dryad (https://doi.org/10.5061/dryad.j2r53 and https://doi.org/10.5061/dryad.sqv9s4n29). BCI trait and seedling dynamics data have been partially made available in other publications and will be posted in Dryad upon publication.

## SUPPLMENTARY MATERIAL

**Supplementary Table 1.**
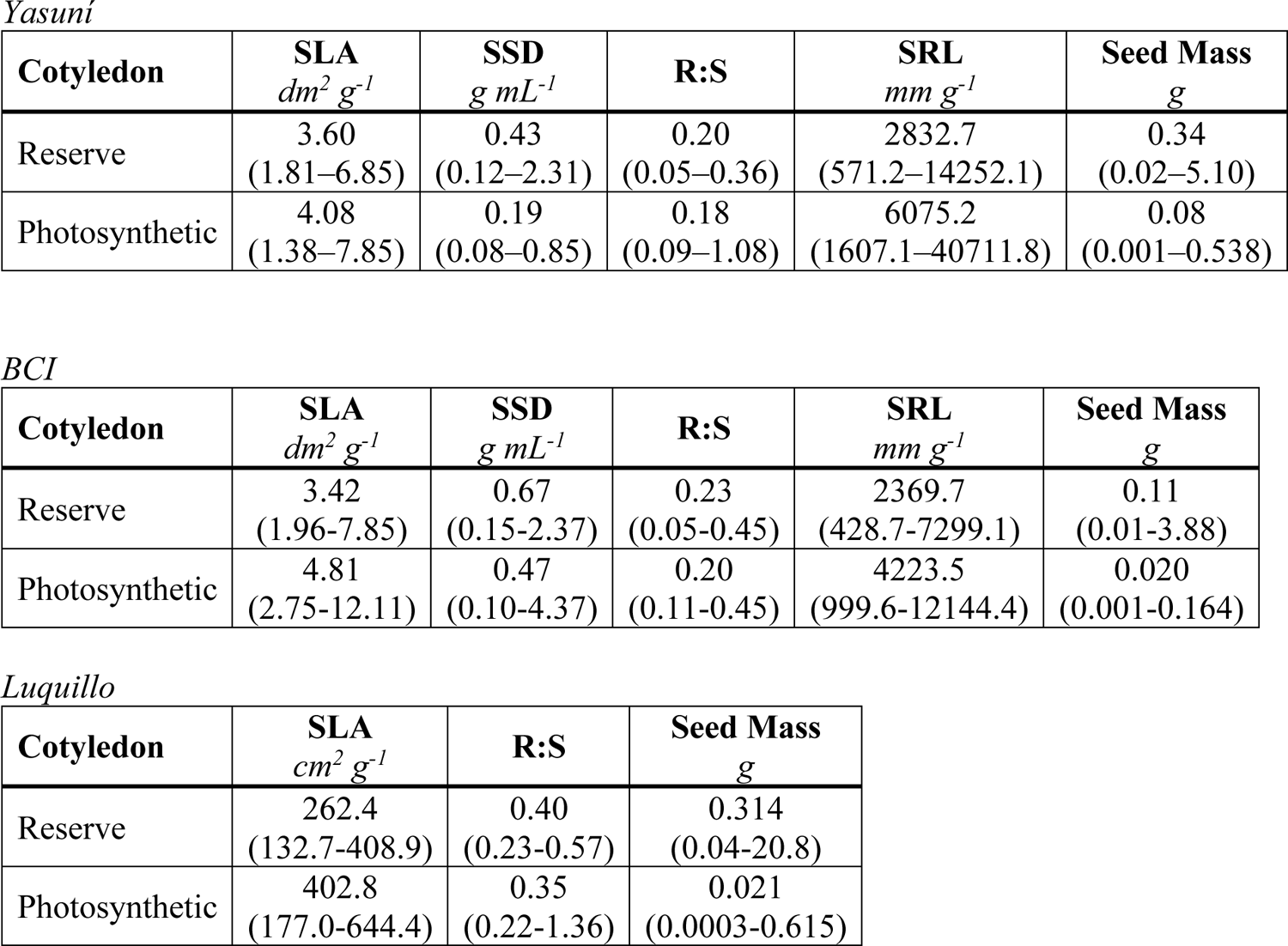
Median trait values (2.5^th^ – 97.5^th^ quantile) of specific leaf area (SLA), stem density (SSD), root:shoot biomass ratio (R:S), specific root length (SRL) and seed mass by cotyledon strategy for species at each site.

**Supplementary Table 2.**
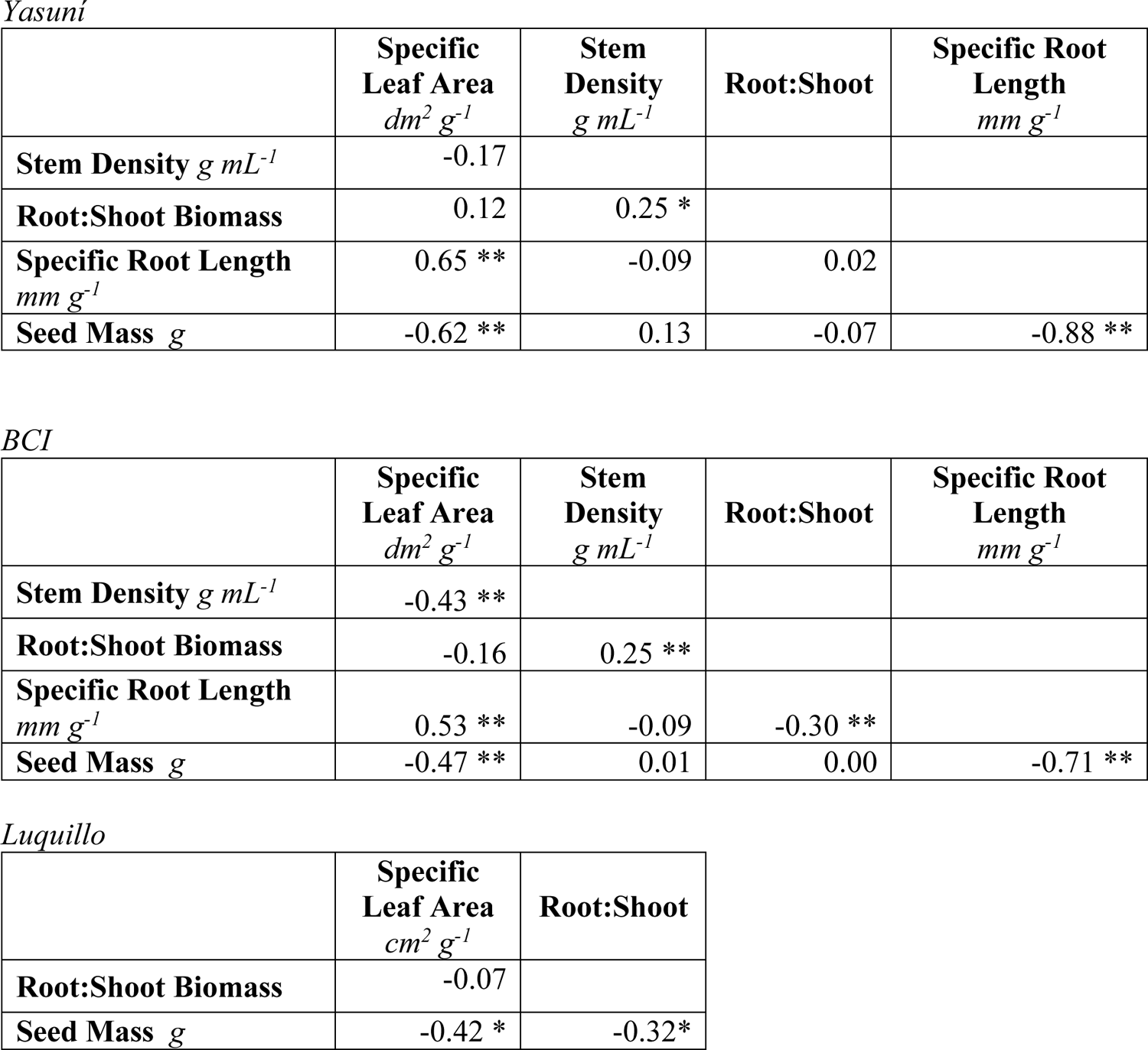
Pairwise Spearman’s *rho* rank correlation coefficients for seedling functional traits and seed mass measured on 84 species at Yasuní, 128 at BCI, and 41 at Luquillo. *p≤0.05, **p≤0.01 after Holm (1979) correction for multiple comparisons.

**Supplementary Table 3.**
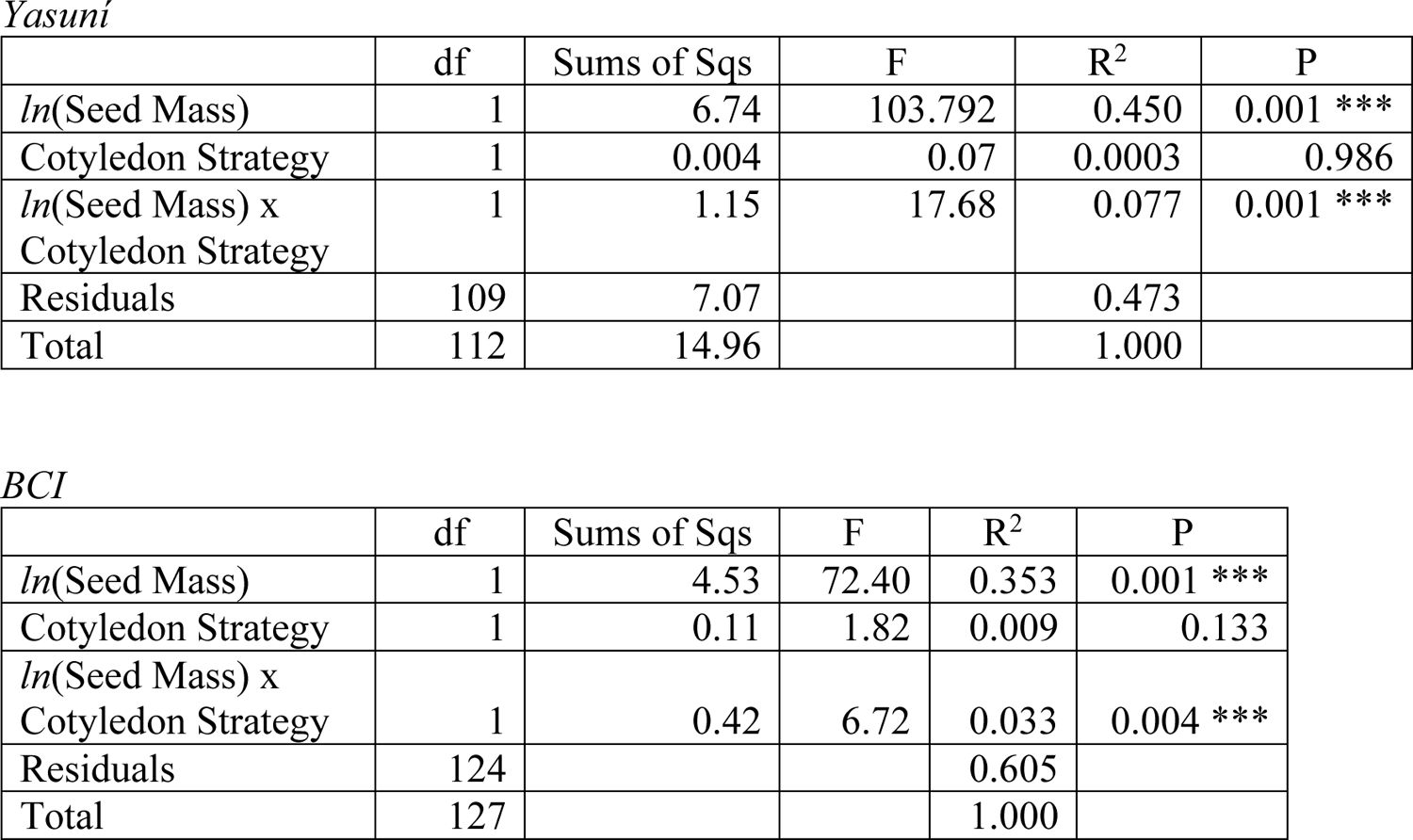
ADONIS results of functional traits NMDS to examine the utility of seed mass or cotyledon strategy as an indicator of seedling traits. Terms are evaluated sequentially, so the analysis considers whether seed mass, followed by cotyledon strategy and its interaction with seed mass, are significant predictors of multivariate variation in leaf, stem, and root traits at Yasuní and BCI.

**Supplementary Table 4.**
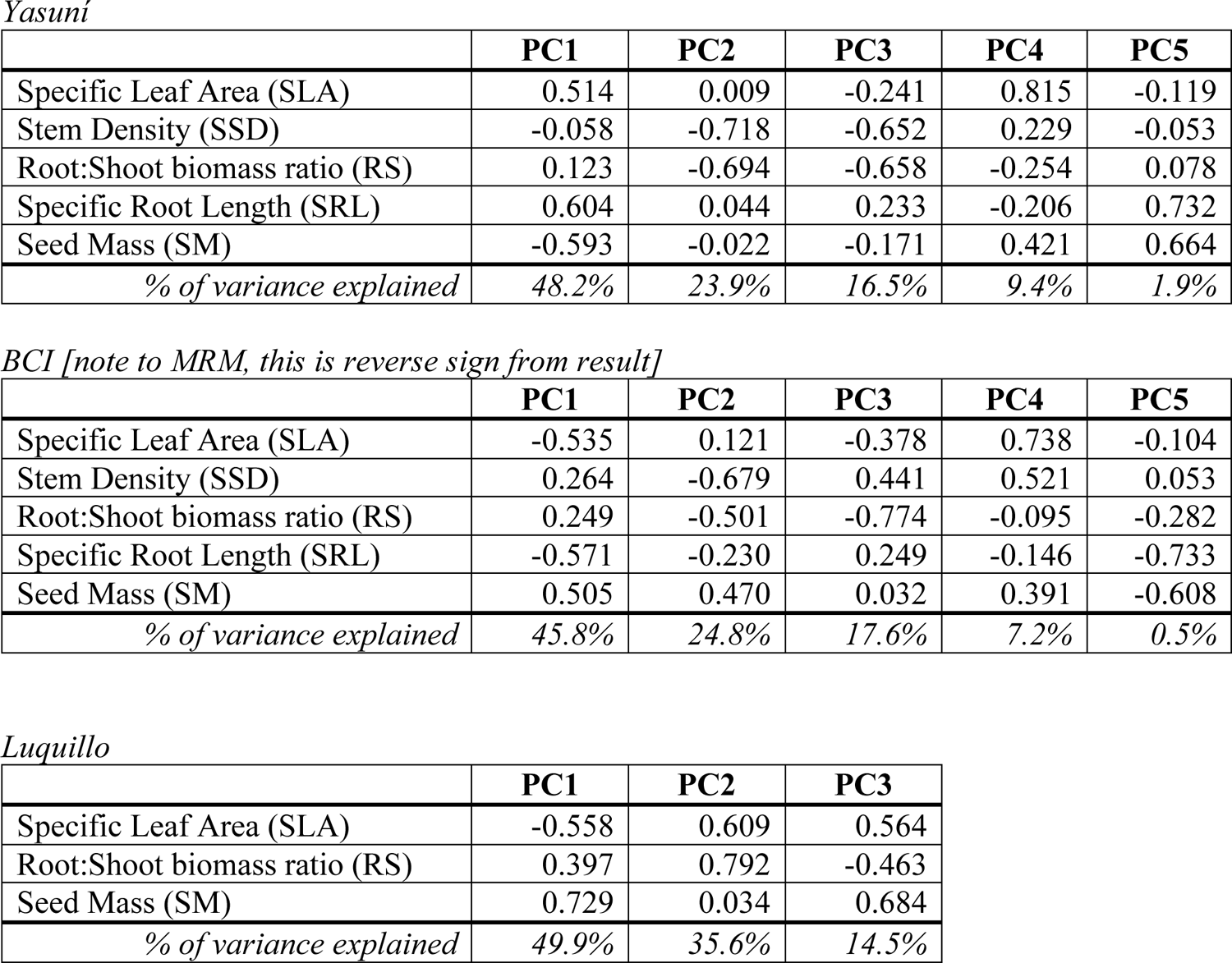
Loadings and percent variance explained for each axis of the PCA on seed mass and seedling functional traits at each site.

**Supplementary Table 5.**
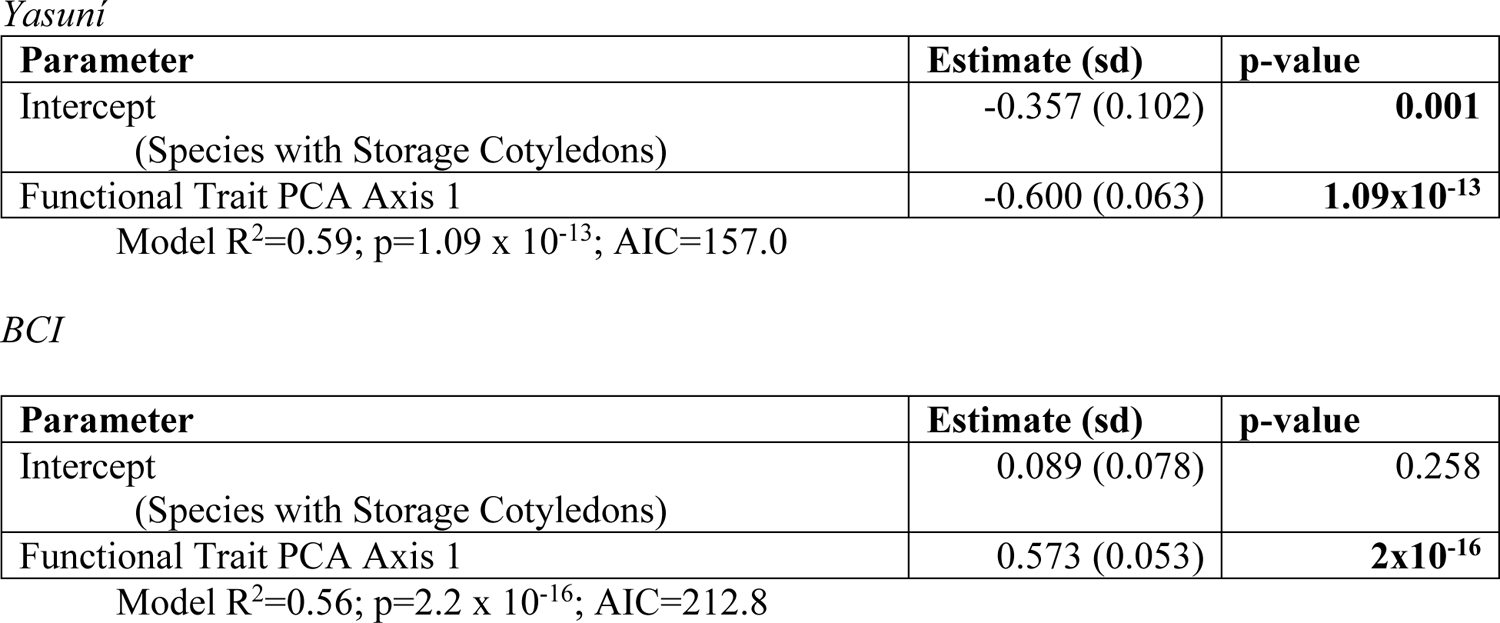
Regression coefficients for simpler model of position along the growth-survival trade-off axis (dependent variable) against the composite functional trait axis (independent variables) without the further explanatory power of the cotyledon type. Here the trait PCA axis includes seed mass in the composite variable. Positive slope estimates indicate a positive relationship with seedling survival and a negative relationship with seedling growth in the first year after recruitment.

## SUPPLMENTARY FIGURE LEGENDS

**Supplementary Figure 1.**
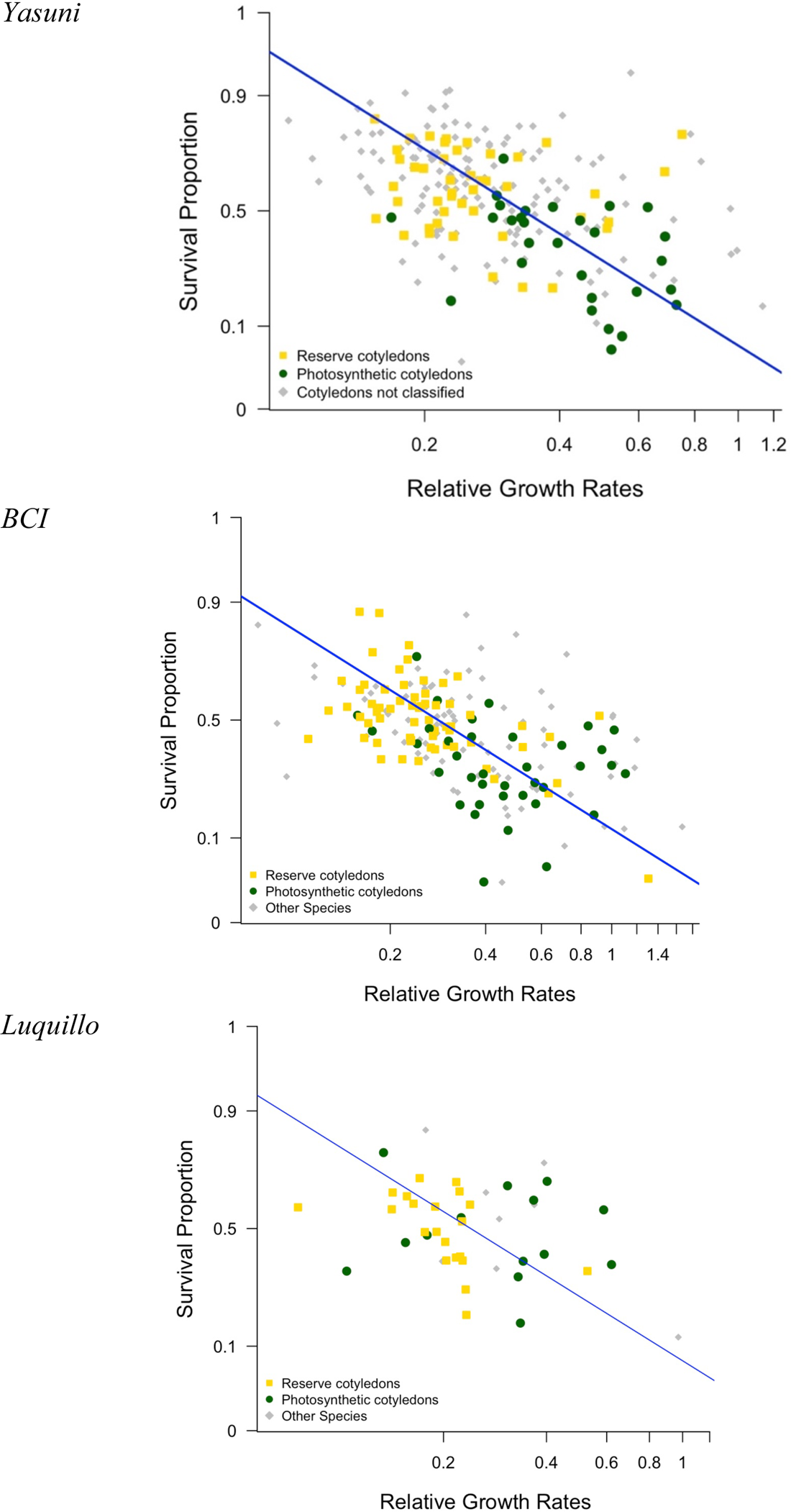
Single-major axis regression of logit-transformed proportion of seedlings that survived their first year and log-transformed average relative growth rate of surviving seedlings. Green and yellow symbols represent species with photosynthetic versus reserve cotyledons, respectively. Gray symbols represent species …

**Supplementary Figure 2.**
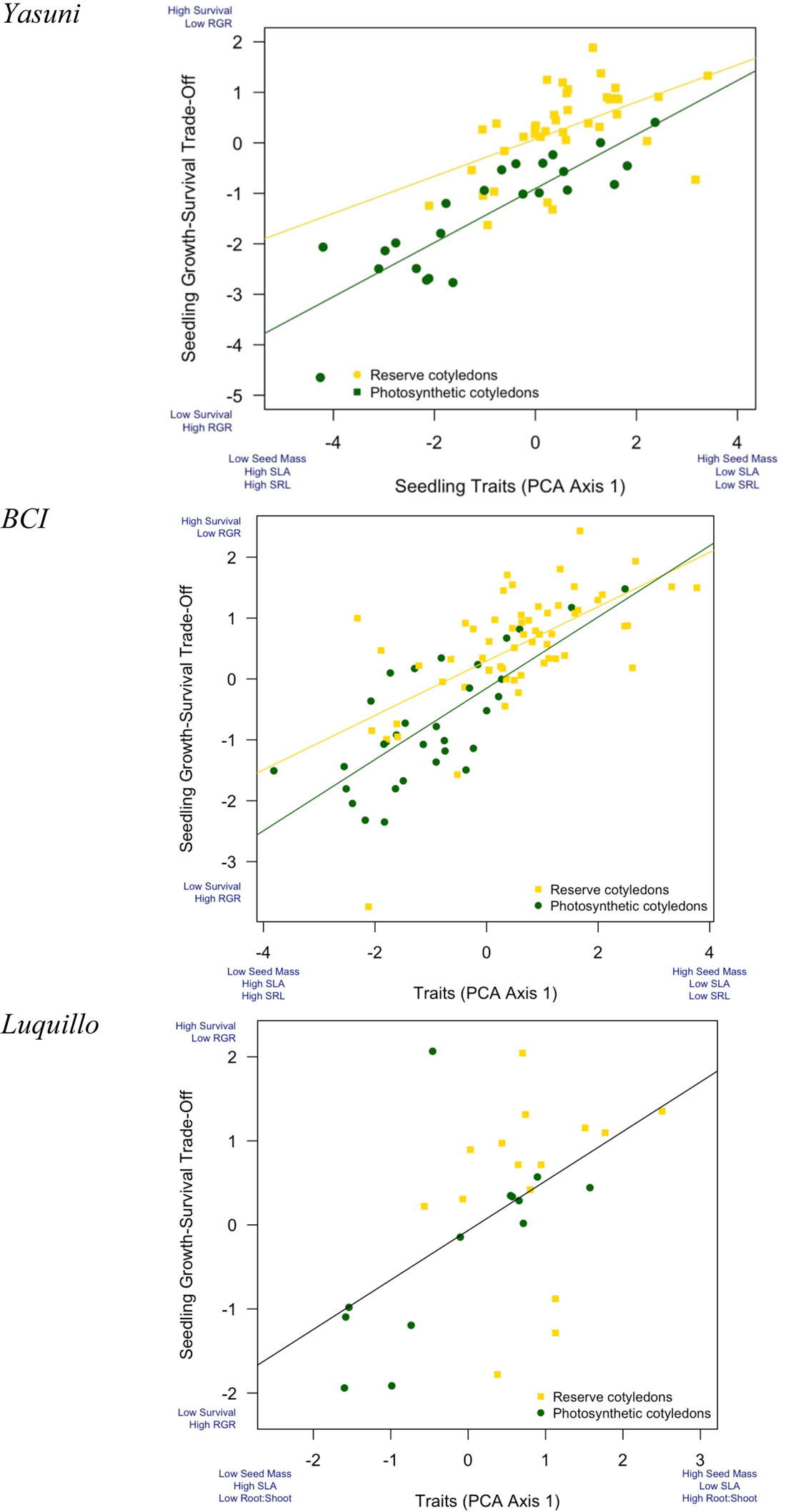
Variation in seedling functional traits (of leaves, stems, roots and seeds) in combination with cotyledon type explains much of the variation in species’ position along a continuum of high survival and high relative growth. Lines are plotted to reflect the model results in Table 1, with separate lines for each cotyledon strategy at Yasuní and BCI.

## Notes

### Competing Interest Statement

The authors have declared no competing interest.

