## Supplementary Materials for "Functional traits of young seedlings predict trade-offs in seedling performance in three neotropical forests"

### SUPPLMENTARY MATERIAL

**Supplementary Table 1.** Median trait values (2.5$^{th}$ – 97.5$^{th}$ quantile) of specific leaf area (SLA), stem density (SSD), root:shoot biomass ratio (R:S), specific root length (SRL) and seed mass by cotyledon strategy for species at each site.

*Yasuní*

| **Cotyledon** | **SLA** *$dm^2\ g^{-1}$* | **SSD** *$g\ mL^{-1}$* | **R:S** | **SRL** *$mm\ g^{-1}$* | **Seed Mass** *g* |
|---|---|---|---|---|---|
| Reserve | 3.60 (1.81–6.85) | 0.43 (0.12–2.31) | 0.20 (0.05–0.36) | 2832.7 (571.2–14252.1) | 0.34 (0.02–5.10) |
| Photosynthetic | 4.08 (1.38–7.85) | 0.19 (0.08–0.85) | 0.18 (0.09–1.08) | 6075.2 (1607.1–40711.8) | 0.08 (0.001–0.538) |

*BCI*

| **Cotyledon** | **SLA** *$dm^2\ g^{-1}$* | **SSD** *$g\ mL^{-1}$* | **R:S** | **SRL** *$mm\ g^{-1}$* | **Seed Mass** *g* |
|---|---|---|---|---|---|
| Reserve | 3.42 (1.96-7.85) | 0.67 (0.15-2.37) | 0.23 (0.05-0.45) | 2369.7 (428.7-7299.1) | 0.11 (0.01-3.88) |
| Photosynthetic | 4.81 (2.75-12.11) | 0.47 (0.10-4.37) | 0.20 (0.11-0.45) | 4223.5 (999.6-12144.4) | 0.020 (0.001-0.164) |

*Luquillo*

| **Cotyledon** | **SLA** *$cm^2\ g^{-1}$* | **R:S** | **Seed Mass** *g* |
|---|---|---|---|
| Reserve | 262.4 (132.7-408.9) | 0.40 (0.23-0.57) | 0.314 (0.04-20.8) |
| Photosynthetic | 402.8 (177.0-644.4) | 0.35 (0.22-1.36) | 0.021 (0.0003-0.615) |

**Supplementary Table 2.** Pairwise Spearman's *rho* rank correlation coefficients for seedling functional traits and seed mass measured on 84 species at Yasuní, 128 at BCI, and 41 at Luquillo. *p≤0.05, **p≤0.01 after Holm (1979) correction for multiple comparisons.

*Yasuní*

| | **Specific Leaf Area** $dm^2\ g^{-1}$ | **Stem Density** $g\ mL^{-1}$ | **Root:Shoot** | **Specific Root Length** $mm\ g^{-1}$ |
|---|---|---|---|---|
| **Stem Density** $g\ mL^{-1}$ | -0.17 | | | |
| **Root:Shoot Biomass** | 0.12 | 0.25 * | | |
| **Specific Root Length** $mm\ g^{-1}$ | 0.65 ** | -0.09 | 0.02 | |
| **Seed Mass** *g* | -0.62 ** | 0.13 | -0.07 | -0.88 ** |

*BCI*

| | **Specific Leaf Area** $dm^2\ g^{-1}$ | **Stem Density** $g\ mL^{-1}$ | **Root:Shoot** | **Specific Root Length** $mm\ g^{-1}$ |
|---|---|---|---|---|
| **Stem Density** $g\ mL^{-1}$ | -0.43 ** | | | |
| **Root:Shoot Biomass** | -0.16 | 0.25 ** | | |
| **Specific Root Length** $mm\ g^{-1}$ | 0.53 ** | -0.09 | -0.30 ** | |
| **Seed Mass** *g* | -0.47 ** | 0.01 | 0.00 | -0.71 ** |

*Luquillo*

| | **Specific Leaf Area** $cm^2\ g^{-1}$ | **Root:Shoot** |
|---|---|---|
| **Root:Shoot Biomass** | -0.07 | |
| **Seed Mass** *g* | -0.42 * | -0.32* |

**Supplementary Table 3.** ADONIS results of functional traits NMDS to examine the utility of seed mass or cotyledon strategy as an indicator of seedling traits. Terms are evaluated sequentially, so the analysis considers whether seed mass, followed by cotyledon strategy and its interaction with seed mass, are significant predictors of multivariate variation in leaf, stem, and root traits at Yasuní and BCI.

*Yasuní*

| | df | Sums of Sqs | F | $R^2$ | P |
|---|---|---|---|---|---|
| *ln*(Seed Mass) | 1 | 6.74 | 103.792 | 0.450 | 0.001 *** |
| Cotyledon Strategy | 1 | 0.004 | 0.07 | 0.0003 | 0.986 |
| *ln*(Seed Mass) x Cotyledon Strategy | 1 | 1.15 | 17.68 | 0.077 | 0.001 *** |
| Residuals | 109 | 7.07 | | 0.473 | |
| Total | 112 | 14.96 | | 1.000 | |

*BCI*

| | df | Sums of Sqs | F | $R^2$ | P |
|---|---|---|---|---|---|
| *ln*(Seed Mass) | 1 | 4.53 | 72.40 | 0.353 | 0.001 *** |
| Cotyledon Strategy | 1 | 0.11 | 1.82 | 0.009 | 0.133 |
| *ln*(Seed Mass) x Cotyledon Strategy | 1 | 0.42 | 6.72 | 0.033 | 0.004 *** |
| Residuals | 124 | | | 0.605 | |
| Total | 127 | | | 1.000 | |

**Supplementary Table 4.** Loadings and percent variance explained for each axis of the PCA on seed mass and seedling functional traits at each site.

*Yasuní*

| | PC1 | PC2 | PC3 | PC4 | PC5 |
|---|---|---|---|---|---|
| Specific Leaf Area (SLA) | 0.514 | 0.009 | -0.241 | 0.815 | -0.119 |
| Stem Density (SSD) | -0.058 | -0.718 | -0.652 | 0.229 | -0.053 |
| Root:Shoot biomass ratio (RS) | 0.123 | -0.694 | -0.658 | -0.254 | 0.078 |
| Specific Root Length (SRL) | 0.604 | 0.044 | 0.233 | -0.206 | 0.732 |
| Seed Mass (SM) | -0.593 | -0.022 | -0.171 | 0.421 | 0.664 |
| *% of variance explained* | *48.2%* | *23.9%* | *16.5%* | *9.4%* | *1.9%* |

*BCI [note to MRM, this is reverse sign from result]*

| | PC1 | PC2 | PC3 | PC4 | PC5 |
|---|---|---|---|---|---|
| Specific Leaf Area (SLA) | -0.535 | 0.121 | -0.378 | 0.738 | -0.104 |
| Stem Density (SSD) | 0.264 | -0.679 | 0.441 | 0.521 | 0.053 |
| Root:Shoot biomass ratio (RS) | 0.249 | -0.501 | -0.774 | -0.095 | -0.282 |
| Specific Root Length (SRL) | -0.571 | -0.230 | 0.249 | -0.146 | -0.733 |
| Seed Mass (SM) | 0.505 | 0.470 | 0.032 | 0.391 | -0.608 |
| *% of variance explained* | *45.8%* | *24.8%* | *17.6%* | *7.2%* | *0.5%* |

*Luquillo*

| | PC1 | PC2 | PC3 |
|---|---|---|---|
| Specific Leaf Area (SLA) | -0.558 | 0.609 | 0.564 |
| Root:Shoot biomass ratio (RS) | 0.397 | 0.792 | -0.463 |
| Seed Mass (SM) | 0.729 | 0.034 | 0.684 |
| *% of variance explained* | *49.9%* | *35.6%* | *14.5%* |

**Supplementary Table 5.** Regression coefficients for simpler model of position along the growth-survival trade-off axis (dependent variable) against the composite functional trait axis (independent variables) without the further explanatory power of the cotyledon type. Here the trait PCA axis includes seed mass in the composite variable. Positive slope estimates indicate a positive relationship with seedling survival and a negative relationship with seedling growth in the first year after recruitment.

*Yasuní*

| **Parameter** | **Estimate (sd)** | **p-value** |
|---|---|---|
| Intercept<br>(Species with Storage Cotyledons) | -0.357 (0.102) | **0.001** |
| Functional Trait PCA Axis 1 | -0.600 (0.063) | **$1.09 \times 10^{-13}$** |

Model $R^2$=0.59; p=$1.09 \times 10^{-13}$; AIC=157.0

*BCI*

| **Parameter** | **Estimate (sd)** | **p-value** |
|---|---|---|
| Intercept<br>(Species with Storage Cotyledons) | 0.089 (0.078) | 0.258 |
| Functional Trait PCA Axis 1 | 0.573 (0.053) | **$2 \times 10^{-16}$** |

Model $R^2$=0.56; p=$2.2 \times 10^{-16}$; AIC=212.8

**SUPPLMENTARY FIGURE LEGENDS**

**Supplementary Figure 1.** Single-major axis regression of logit-transformed proportion of seedlings that survived their first year and log-transformed average relative growth rate of surviving seedlings. Green and yellow symbols represent species with photosynthetic versus reserve cotyledons, respectively. Gray symbols represent species …

**Supplementary Figure 2.** Variation in seedling functional traits (of leaves, stems, roots and seeds) in combination with cotyledon type explains much of the variation in species' position along a continuum of high survival and high relative growth. Lines are plotted to reflect the model results in Table 1, with separate lines for each cotyledon strategy at Yasuní and BCI.

**Supplementary Figure 1**

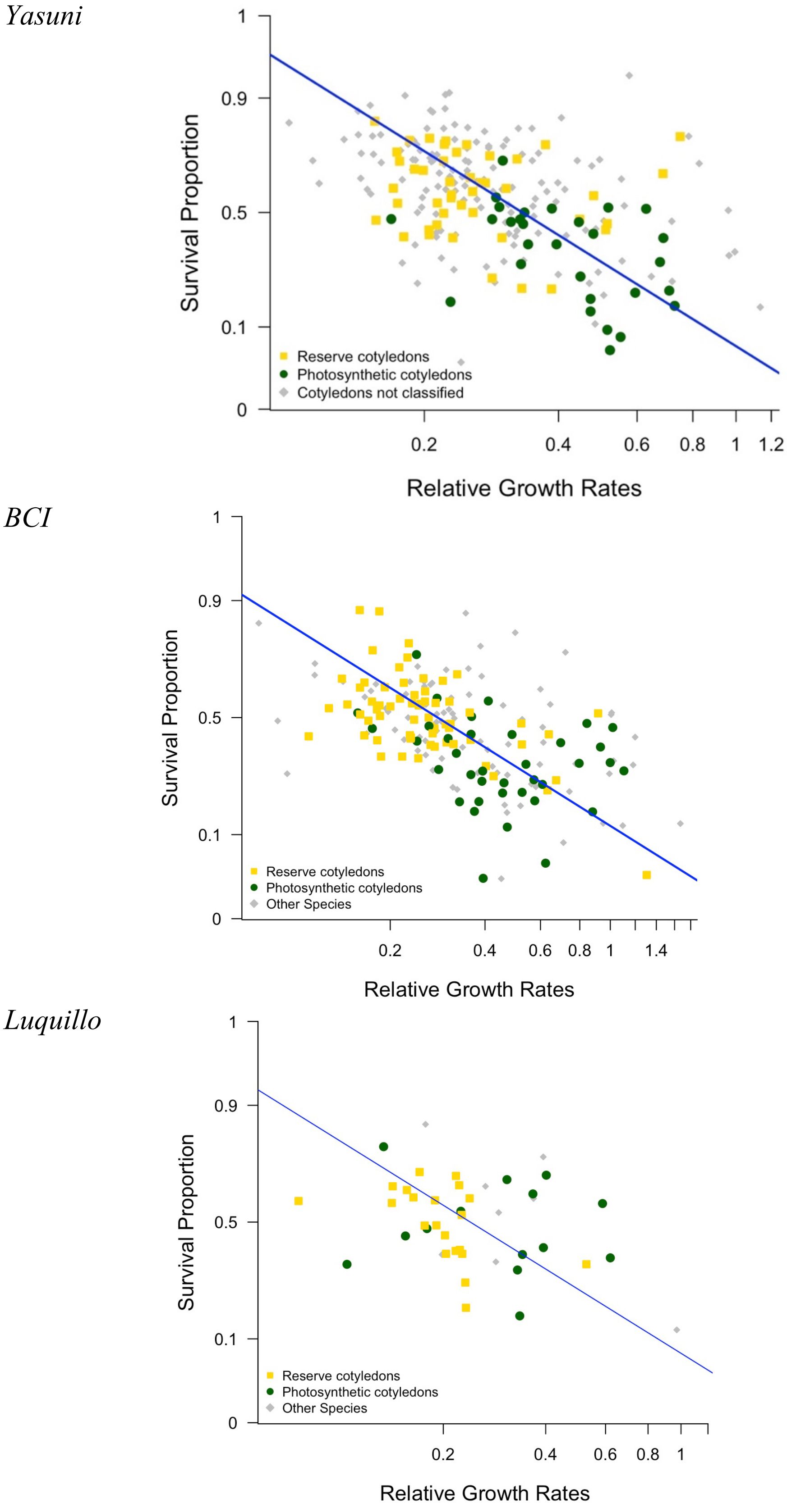

**Supplementary Figure 2**

*Yasuni*

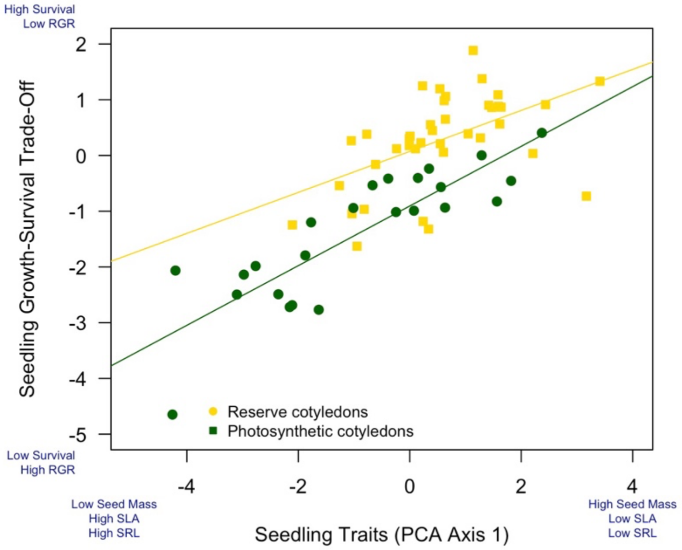


*BCI*

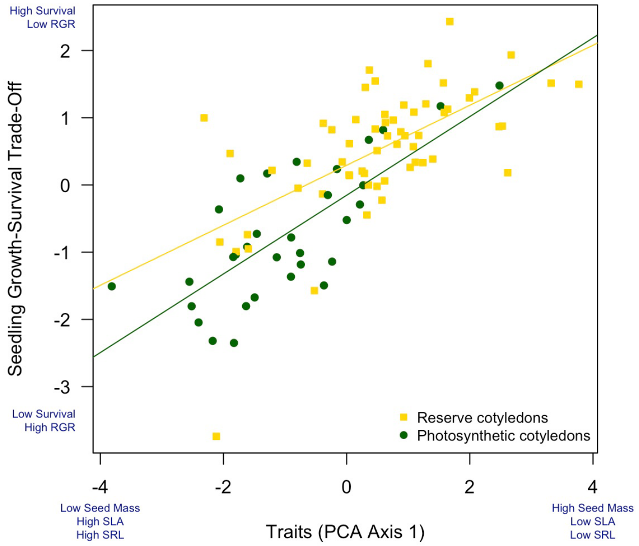


*Luquillo*

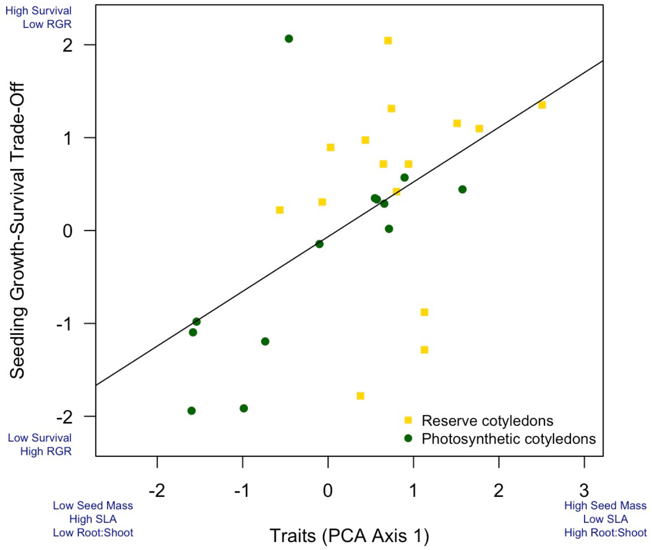
